# Sexually dimorphic gene expression and transcriptome evolution provides mixed evidence for a fast-Z effect in *Heliconius*

**DOI:** 10.1101/380030

**Authors:** A Pinharanda, M Rousselle, SH Martin, JJ Hanly, JW Davey, S Kumar, N Galtier, CD Jiggins

## Abstract

Sex chromosomes have different evolutionary properties as compared to the autosomes due to their hemizygous nature. In particular, recessive mutations are more readily exposed to selection, which can lead to faster rates of molecular evolution. Here, we report patterns of gene expression and molecular evolution in the sex chromosomes of a group of tropical butterflies. We first improved the completeness of the *Heliconius melpomene* reference annotation, a neotropical butterfly with a ZW sex determination system. Then we sequenced RNA from male and female whole abdomens and female ovary and gut tissue to identify sex and tissue specific gene expression profiles in *H. melpomene*. Using these expression profiles we compare sequence divergence and polymorphism, the strength of positive and negative selection and rates of adaptive evolution for Z and autosomal genes between two species of *Heliconius* butterflies, *H. melpomene* and *H. erato*.

We show that the rate of adaptive substitutions is higher for Z as compared to autosomal genes, but contrary to expectation it is also higher for male as compared to female biased genes. There is therefore mixed evidence that hemizygosity influences the rate of adaptive substitutions. Additionally, we find no significant increase in the rate of adaptive evolution or purifying selection on genes expressed in ovary tissue, a heterogametic specific tissue. Together our results provide limited support for fast-Z evolution. This contributes to a growing body of literature from other ZW systems that also provide mixed evidence for a fast-Z effect.

## Introduction

Heteromorphic sex chromosomes have different evolutionary properties compared to autosomes (Rice, 1984). Specifically, because recessive mutations are exposed to selection more readily on the sex chromosomes, positive and purifying selection - as well as the strength of genetic drift - are expected to result in different rates of molecular evolution between sex chromosomes and autosomes. An increased evolutionary rate of sex chromosomes relative to autosomes, known as the fast-X effect (Charlesworth *et al*., 1987), has been observed in *Drosophila* (Ávila *et al*., 2014). X genes are expected to diverge faster between species than autosomal genes mainly due to the higher substitution rate of recessive, advantageous mutations. However, this process is also influenced by: 1) patterns of selection in males versus females; 2) mutation; 3) recombination and 4) demography (Orr and Betancourt, 2001; Kirkpatrick and Hall, 2004; Vicoso and Charlesworth, 2006; 2009; Pool and Nielsen, 2007; Orr, 2010; Connallon *et al*., 2012).

Patterns of molecular evolution on the sex chromosomes are particularly influenced by gene expression patterns. Sexually dimorphic expression is often caused by natural and/or sexual selection favouring phenotypes that influence the fitness of one of the sexes (Grath and Parsch, 2016). In species with genetic sex determination the majority of sexually dimorphic traits result from the differential expression of genes present in both male and female genomes (Ellegren and Parsch, 2007). Sex biased expression is common across taxa from mammals (Rinn and Snyder, 2005) to Diptera (Assis *et al*., 2012), reptiles (Cox *et al*., 2017), birds (Mank *et al*., 2010) and Lepidoptera (Rousselle *et al*., 2016). For example, in *Drosophila melanogaster*, 57% of genes have been categorised as sex biased (Assis *et al*., 2012); and, in *Heliconius melpomene*, analysis of two different tissues identified up to 29% of expressed genes as sex biased (Walters *et al*., 2015). The vast majority of genes that exhibit sexually dimorphic expression are active in reproductive tissues and tend to also have distinctive rates of molecular evolution compared to genes without dimorphic expression (Parisi *et al*., 2003; 2004; Avila *et al*., 2015). Ultimately, the identification of sex biased genes and subsequent analysis of patterns of molecular evolution will contribute to a better understanding of the evolutionary forces shaping sex chromosome and autosome evolution (Kirkpatrick and Hall, 2004; Zhang *et al*., 2004; Assis *et al*., 2012).

Empirical studies of the fast-X effect typically measure two different metrics: 1) the ratio of non-synonymous to synonymous substitution rates (dN/dS); and 2) the amount of adaptive evolution (α) using the McDonald-Kreitman (MK) test (McDonald and Kreitman, 1991). Studies measuring dN/dS usually test for “faster-X divergence”. Although this approach may be useful for comparing sex chromosome and autosomal divergence measuring only the relative rate of non-synonymous substitutions captures the effects of both adaptive and neutral (or slightly deleterious) mutations. Estimates of α can better test for an excess of adaptive substitution in the sex chromosome (“faster-X adaptation”) by combining measures of within-species polymorphism and between-species divergence, but α is still sensitive to the rate of accumulation of slightly deleterious mutations and demography (Fay 2011). For instance, an increase in N_e_ is expected to result in decreased dN/dS and increased α even when the rate of adaptive substitutions remains unchanged. To overcome this problem extensions of the MK test such as ω_a_ were developed to estimate the rate of adaptation by calculating the frequency distribution of polymorphism after correcting for demographic history and distribution of deleterious effects at functional sites (Galtier, 2016).

The analysis of evolutionary rates between sex and autosomal genes, however, has produced mixed evidence in support of fast-X evolution (Meisel and Connallon, 2013). In some taxa there is strong evidence for faster-X divergence but not faster-X adaptation or vice versa (Meisel and Connallon, 2013). For example, the first calculations of faster-X divergence were carried out in *Drosophila* where support for elevated dN/dS in X genes has been mixed.

Studies that used autosome-to-X translocations to control for gene content effect did not reach a consensus on the existence of faster-X divergence (Counterman *et al*. 2004; Thornton *et al*., 2006; Zhou and Bachtrog, 2012) but X-linked duplicate genes have elevated dN/dS compared to autosomal duplicates (Thornton and Long, 2002). Signals of faster-X sequence divergence in *Drosophila* have been shown to affect non-coding regulatory regions as well, and might be at least partly explained by differences in gene composition on the X versus the autosomes (Hu *et al*., 2013). However, faster-X divergence in other taxa has received stronger support. For example in humans, chimpanzees and rodents dN/dS is higher for X genes (Nielsen *et al*., 2005; Mank, Vicoso, *et al*., 2010).

In contrast, whole-genome analyses of adaptive substitutions have resulted in stronger evidence for faster-X adaptation in *Drosophila* (Mackay *et al*., 2012), while support for faster-X adaptation in vertebrates is less clear. McDonald-Kreitman tests support faster-X adaptation in wild mouse populations (Baines and Harr, 2007) but, for the European rabbit *(Oryctolagus cuniculus)*, a clear faster-X adaptation signal is only present in populations with large effective population sizes (Carneiro *et al*., 2012).

Taxa with ZW sex determination provide an interesting contrast. For female-heterogametic taxa such as birds, females only have one copy of the Z chromosome. A fast-Z effect may be expected to result from the expression of recessive mutations on the Z chromosome as Z genes are immediately exposed to selection in females (Charlesworth *et al*., 1987). In birds, fast-Z divergence has been reported, but Z male biased genes were not less accelerated than unbiased genes or female biased genes (Wright *et al*., 2015). This would not be expected if the fast-Z effect was driven by recessive beneficial mutations, and so it was suggested that fast-Z in birds does not reflect positive selection (Mank, Nam, *et al*., 2010; Wright *et al*., 2015).

For Lepidoptera results have also been mixed. Sackton *et al*. (2014) reported that faster-Z evolution was driven by position selection in silkworms. But, in satyrine butterflies, there were no significant differences in adaptive evolutionary rates between the Z and the autosomes (no fast-Z adaptation). However, the comparison of male biased, female biased and unbiased Z genes in satyrine butterflies revealed increased purifying selection against recessive deleterious mutations in female biased Z genes (Rousselle *et al*., 2016). Considerable uncertainty therefore remains regarding the prevalence and magnitude of the fast-X/Z effect on divergence and adaptation.

Here we investigate the effects of hemizygosity on the rates of adaptive substitution in the neotropical butterfly genus *Heliconius*, a ZW sex determination system, by analysing polymorphism, divergence and gene expression genome-wide. We test whether there is a fast-Z effect in *Heliconius* using two species from the *H. melpomene* and *H. erato* clades which diverged 13 million years ago (synonymous divergence = 0.16) (Kozak *et al*., 2015; Martin *et al*., 2016). Previous analyses of *Heliconius* transcriptome data have focused on the evolution of dosage compensation and the impact of sex specific dosage on the levels of gene expression (Walters *et al*., 2015). In this study, using the same transcriptome data, we first compute sex biased expression. Then, accounting for sex biased gene expression, we: 1) calculate coding sequence divergence and polymorphism in *H. melpomene;* and 2) assess the strength of positive and negative selection, and rates of adaptive evolution between *H. melpomene* and *H. erato*. We then analyse newly generated female transcriptome data from *H. melpomene* ovary and gut tissue in order to investigate whether genes expressed in the reproductive tissue of the heterogametic sex have higher rates of adaptive evolution than those expressed in somatic tissues.

## Material and Methods

### Updated *H. melpomene* annotation

The Hmel2 annotation of the *H. melpomene* genome has 13 178 predicted transcripts spanning 16 897 139 bp (The Heliconius Genome Consortium, 2012; Davey et al., 2016). The Hmel2 annotation is incomplete, as there are 20 118 high quality predicted transcripts in *H. erato* spanning 33 669 374 bp (van Belleghem et al., 2017). To improve the completeness of the annotation for *H. melpomene* we downloaded RNA-seq reads from NCBI repositories ArrayExpress ID: E-TAB-1500 (Briscoe et al., 2013), and BioProject PRJNA283415 (Walters et al., 2015), published since Hmel1 release. We also used data from 10 wing RNA-seq libraries (Hanly 2017). We used the BRAKER1 pipeline to perform unsupervised RNA-seq based genome annotation (Hoff *et al*., 2016). GeneMark-ET was used to perform iterative training, generating initial gene structures and AUGUSTUS was used for training and subsequent integration of RNA-seq read information into the final gene predictions (Stanke et al., 2008; Lomsadze et al., 2014; Hoff et al., 2016). This resulted in 26,017 predicted transcripts spanning 32,222,367 bp. 6,532 of these transcripts were considered repeat proteins based on 90% single hit match to repeat databases Repbase were removed (Bao *et al*., 2015). We transferred 428 manually annotated genes (441 transcripts/protein) from the original Hmel2 annotation and removed any BRAKER1 predictions that overlapped. We also transferred 189 genes (189 transcripts/proteins) that have been manually annotated and published since Hmel2 release. Specifically, we transferred 73 gustatory receptors; 31 immune response and 85 Glutathione-S-transferases and Glucuronosyltransferases (Briscoe et al., 2013; van Schooten et al., 2016; Yu et al., 2016) and removed any BRAKER1 predictions that were overlapping. Moreover, BRAKER1 predictions that had 1-to-1 overlaps with Hmel2 names were replaced by their original Hmel2 name. For many-to-1 mapping between the BRAKER1 predictions and Hmel2, Hmel2 names were reused and a suffix of g1/g2/g3/etc was added. The rest were renamed from HMEL030000 onwards.

### Samples for gene expression analysis

Gene expression data was calculated using: 1) Illumina 100bp paired-end RNA-seq data from 5 Panamanian *H. m. rosina* whole-male abdomens, and 5 Panamanian *H. m. rosina* whole-female abdomens, downloaded from GenBank (BioProject PRJNA283415) (Walters *et al*., 2015); and 2) newly sequenced Illumina HiSeq 2500 150bp paired-end directional (stranded) RNA-seq data from ovary tissue of 7 young (1h) and 6 old (20 days) *H. m. rosina* females, and from gut tissue of 6 young (1h) and 6 old (20 days) *H. m. rosina* females (25 samples from 13 different individuals, Supplementary Table S1).

For these 25 samples *H. m. rosina* females were reared in insectaries in Gamboa, Panama. *Passiflora platyloba* potted plants were monitored daily and 5^th^ instar caterpillars were removed and taken to the laboratory in large individual containers where they were allowed to pupate and emerge at a constant temperature (24-25°C). The pupating containers in the laboratory were monitored several times a day. When a female emerged, it was either: 1) returned to the insectaries to be mated to a *H. m. rosina* male (Treatment: old, Supplementary Table S1); or 2) dissected 1h after eclosion under controlled laboratory conditions (Treatment: young, Supplementary Table S1). Mated females were kept in individual 1m x 1m x 2m cages for 20 days until dissection.

Guts and ovaries were dissected in RNAlater (ThermoFisher, Waltham, MA) at 24-25°C and tissue was stored in RNAlater at 4°C for 24h and −20°C thereafter. Total RNA was extracted with a combined guanidium thiocyanate-phenol-chloroform and silica matrix protocol using TRIzol (Invitrogen, Carlsbad, CA), RNeasy columns (Qiagen, Valencia, CA) and DNaseI (Ambion, Naugatuck, CT). mRNA was isolated from total RNA via poly-A pull-down, and directional cDNA library preparation and sequencing (Illumina HiSeq 2500, 150bp paired end) were performed by Novogene Bioinfomatics Technologies (Hong Kong, China) (Supplementary Table S1).

### Read mapping, counting and identification of sex and ovary and gut biased genes

FASTQ reads were aligned to gene sequences from *H. melpomene* v2.5 annotation using HISAT2 (Kim *et al*., 2015) with default mapping parameters. Mapping statistics were calculated using samtools flagstat (v1.2) (Li *et al*., 2009). We used htseq-count to determine the number of aligned sequencing reads mapped to each genic feature (HTSeq v0.6.1; python v2.7.10; option: -m union) (Anders *et al*., 2015).

Estimation of variance-mean dependence from the count data was performed with DESeq2 (v1.14.1) (Love *et al*., 2014) using Bioconductor v3.4 and R v3.2.5, using the constructor function DESeqDataSetFromHTSeqCount(design=~batch+sex) for sex biased genes and DESeqDataSetFromHTSeqCount(design=~batch+tissue) for ovary and gut biased genes. All the result tables were built using the DESeq2 results() function (options: betaPrior=false, test=Wald). We filtered the results as in Walters *et al*. (2015) with FDR < 0.05 (alpha=0.05) (Walters *et al*., 2015). We defined male, female and unbiased genes as in Rousselle *et al*. (2016); male biased genes have log_2_ fold change significance threshold < 0.66 (option: lcfThreshold<0.66), female biased genes have log2 fold change significance threshold > 1.5 (option: lcfThreshold>1.5) and the others were classified as unbiased.

### Extraction of orthologous genes, coding sequence alignment and SNP calling

OrthoFinder was used to identify orthologous groups of genes in the *H. melpomene* and the *H. erato* transcriptomes (options: -t 48 -a 6). 1-1 orthologous gene sequences were selected for use in subsequent analysis (Supplementary Table S2). Using Gff-Ex, a genome feature extraction package (Rastogi and Gupta, 2014), we extracted coding sequences from: 1) 10 whole-genome short-read re-sequenced wild *H. m. rosina* from Panama (Supplementary Table S3; Van Belleghem et al., 2018) mapped to Hmel2 (Davey *et al*., 2016) with bwa-mem (Li and Durbin, 2009); 2) the reference *H. erato* genome (Van Belleghem *et al*., 2017).

For the 10 whole-genome re-sequence *H. m. rosina* samples (van Belleghem *et al*., 2018), genotypes were called using HaplotypeCaller (GATK v3.4-0-g7e26428) (DePristo *et al*., 2011), and genotypes were designated as missing if the read depth for a given individual at a given site was <8. Coding sequences for 1-1 orthologous genes were extracted in fasta format from 1) and 2) and aligned using MACSE, accounting for frameshifts and stop codons (Ranwez *et al*., 2011).

### Calculation of diversity and selection statistics for 1-1 ortholog alignments between *H. melpomene* and *H. erato: Classic approach*

The adaptive substitution rate was estimated by comparing synonymous and non-synonymous variation in the polymorphism and divergence compartments, as first proposed by McDonald & Kreitman, 1991; see also Bustamante *et al*., 2005, and Mcpherson *et al*., 2007). We first used the original MK test (referred to as *Classic approach* hereafter) to estimate the rate of adaptive substitution for all genes found to be orthologous between *H. melpomene* and *H. erato*. We calculated: 1) synonymous polymorphism (P_s_) and 2) non-synonymous polymorphism (P_n_) in *H. melpomene*, as well as 3) synonymous fixed divergence (dS), and 4) non-synonymous fixed divergence (dN) between *H. melpomene* and *H. erato*. We estimated the rate of adaptive molecular evolution (α) between the two species as:

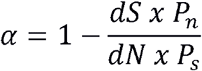

α assumes that non-synonymous mutations are either adaptive, neutral, or strongly deleterious (McDonald and Kreitman, 1991), with −∞ > α ≥ 1, where α = 0 represents the null hypothesis that non-synonymous mutations are neutral (dN/dS = P_n_/P_S_). α > 0 corresponds to dN/dS > P_n_/P_S_ and indicates positive selection, whereas α < 0 cprresponds to dN/dS < P_n_/P_S_ and indicates negative selection. These values were calculated using the EggLib C++ function polymorphismBPP (v2.1.11) (De Mita and Siol, 2012) and Bio++ (v2.2.0) (Dutheil and Boussau, 2008) in python (v2.7.5) using scripts adapted from https://github.com/tatumdmortimer (last accessed 09/04/2018) (O’Neill *et al*., 2015).

### Calculation of diversity and selection statistics for 1-1 ortholog alignments between *H. melpomene* and *H. erato: Modelling approach*

The *Classic approach* to the MK test is robust to differences in mutation rates and variation in coalescent histories across genomic locations (McDonald & Kreitman, 1991). Inference of positive selection using the *Classic approach* of the MK test is not robust, however, to the occurrence of slightly deleterious mutations and demographic change. To account for these confounders, we used a *Modelling* approach to estimate the strength of positive and purifying selection in addition to the *Classic approach* described above, using the method of Eyre-Walker and Keightley (2009) as implemented in Galtier (2016) and Rousselle *et al*. (2016).

The *Modelling approach* uses the frequency distribution of polymorphism to assess the distribution of deleterious mutations at functional sites. This elaborates on the *Classic approach* of the MK test by modelling the distribution of fitness effects (DFE) of deleterious non-synonymous mutations as a negative Gamma distribution. The model is fitted to the synonymous and non-synonymous site frequency spectra (SFS) and the expected dN/dS under near-neutrality is inferred. The difference between the observed and expected dN/dS provides an estimate of the proportion of adaptive non-synonymous substitutions (α). The per mutation rate of adaptive substitutions is calculated as:

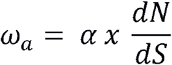

and the per mutation rate of non-adaptive substitutions is calculated as:

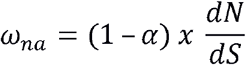

### Gene expression level and π_n_/π_s_ ratios

We calculated reads per kilobase per million (RPKM) as:

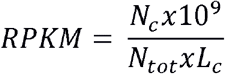

To test whether gene expression level and chromosome type have a significant effect on π_n_/π_s_ ratios we used a multiple regression analysis. We established the linear model:

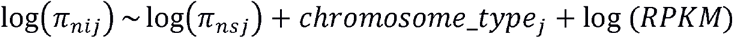

using R (v3.2.5). Where N_c_ is the number of reads mapped to the genic feature, N_tot_ is the total number of reads mapped in the sample, and L_c_ is the length of the genic sequence in base pairs (Mortavazi *et al*., 2008). RPKM_i_ is the mean RPKM of gene *i* across the 10 individuals. Chromosome type is either autosome or sex chromosome as assigned in Hmel2 reference genome (Davey *et al*., 2016). 477 genes with no polymorphism were removed from the analysis. We plotted diagnostic plots of residuals versus fitted values.

## Results

### Hmel2.5 annotation and 1-1 orthologue prediction with *H. erato*

There are 20,118 transcripts predicted in *H. erato* (van Belleghem et al., 2017) and 20,097 genes (21,565 transcripts/proteins) in *H. melpomene* Hmel2.5. OrthoFinder returned 11,062 clusters of genes, 8085 of which included exactly one sequence per species. 14,841 (73.8%) of the total number of genes in *H. erato* were assigned to an orthogroup; and 14,857 (68.6%) of the total number of genes in *H. melpomene* version Hmel2.5 were assigned to an orthogroup (Supplementary Table S2). Conversely, the *H. melpomene* Hmel2 annotation has 13 019 predicted gene models (Davey *et al*., 2016). Using the Hmel2 annotation, OrthoFinder returned 9,320 clusters of genes, 6846 of which included exactly one sequence per species (i.e. single-copy orthogroups). 13 744 (68.3%) genes were assigned to an orthogroup in *H. erato* and 10 530 (80.9%) were assigned to an orthogroup in *H. melpomene*. The Hmel2.5 annotation set for *H. melpomene* is therefore more comparable to the published *H. erato* gene annotation and is more appropriate for future gene-based analysis in *H. melpomene*. The Hmel2.5 annotation has 1,093 genes mapping to the Z chromosome and 18,835 mapping to the autosomes. This new version of *H. melpomene* genome annotation was numbered Hmel2.5 (available at LepBase http://ensembl.lepbase.org/Heliconius_melpomene_melpomene_hmel25/Info/Index, last accessed 20 Jun 2018, Challis *et al*., BioRxiv preprint).

### RNA-sequencing and read mapping

Analysis of gene expression profiles in the data retrieved from Walters *et al*. (2015) by principal component, the first principal component separates gene expression in whole abdomen by sex and explains 97% of variance (Supplementary Figure S1). The 25 *H. melpomene* samples sequenced for this project have a median total number of reads of 34.86 M (min. 27.81 M; max. 46.12 M), similar to previously published gene expression studies in *Heliconius* (Briscoe *et al*., 2013; Walters *et al*., 2015). Mapping success is high compared to other published studies (e.g. Yu et al., 2016 and Walters et al., 2015) (Supplementary Table S1). We analysed data from two different time points and from non-sex and sex-specific tissue separately (Treatment: Young and Old). There is a clear separation of the 25 samples by tissue when we compare gene expression profiles between them. In total, 51% of the total variance is explained by the two first principal components. PC1 separates the samples by tissue and explains 40% of variance. PC2 explains 11% of total variance and separates samples by age (Supplementary Figure S3). Ovarian tissue clusters by age more tightly than non-sex specific tissue (Gut) (Supplementary Figure S4A and S4B).

### Coding sequence divergence does not support a significant fast-Z effect

We first compared rates of Z and autosomal sequence divergence using dN/dS comparisons of 1-1 orthologous genes between *H. melpomene* and *H. erato*. The dN/dS ratio for the Z chromosome genes is not significantly higher than dNdS for autosomal genes (dN/dS_Auto_=0.110; 95% CI=[0.106-0.113]; dN/dS_Z_=0.120; 95% CI= [0.098-0.145]), indicating no obvious faster Z divergence of coding sequence.

More highly expressed genes are more exposed to selection, so in a female heterogametic organism with a fast-Z effect, genes with female-biased expression are expected to have higher rates of amino acid substitution if dN/dS is driven by positive selection. However, the dN/dS ratio of Z female biased genes (dN/dS_Z_=0.120; 95% CI=[0.069-0.183]) was not significantly different to that for male biased genes (dN/dS_Z_=0.148 [0.122-0.172]) or unbiased genes (dNdS_Z_=0.107; 95% CI=[0.078-0.143]). By contrast, among autosomal genes, those that are unbiased have a significantly lower coding sequence divergence compared to both male biased and female biased autosomal genes (dNdS_Z_=0.0978; 95% CI=[0.093-0.102]) (Table 1).

**Table 1.**
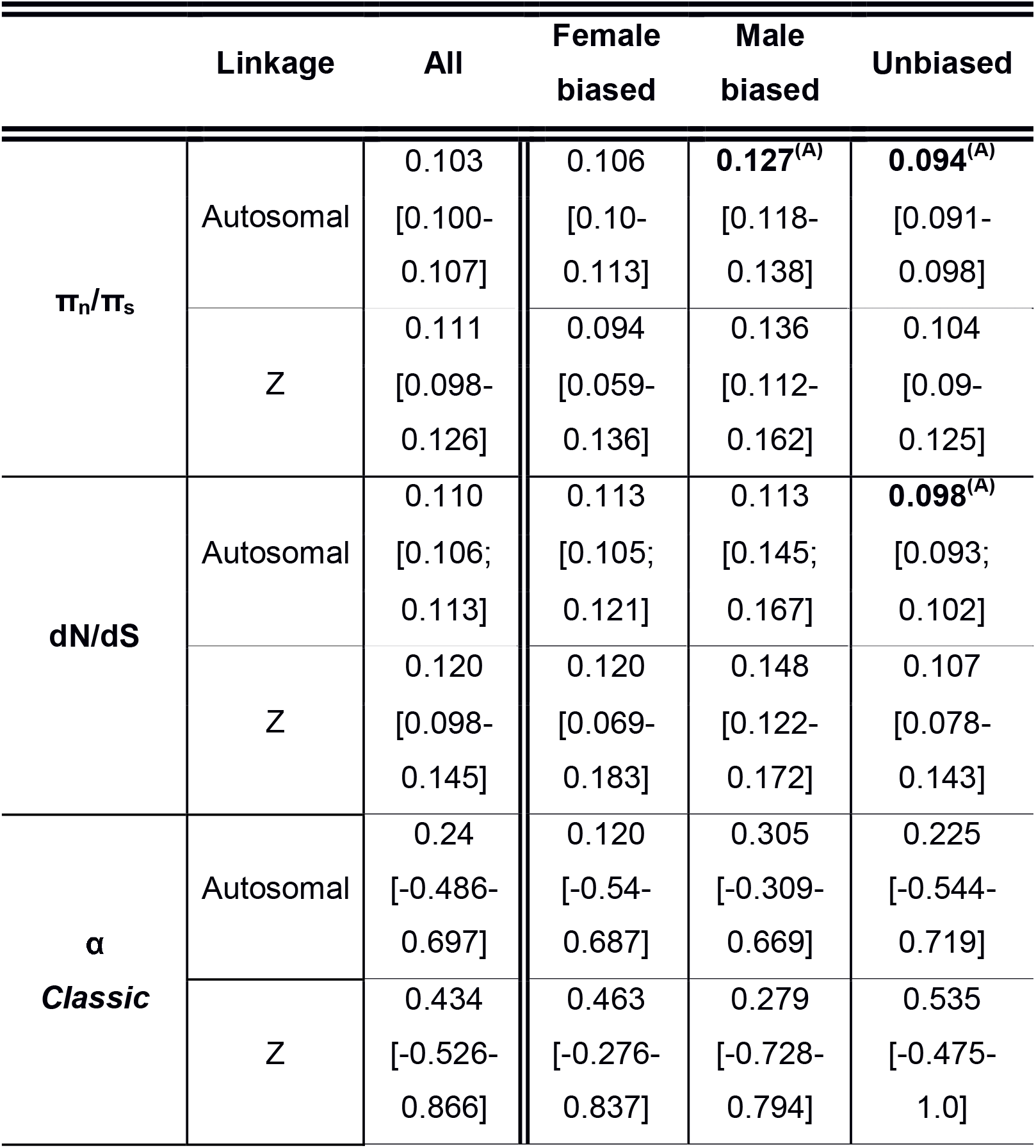
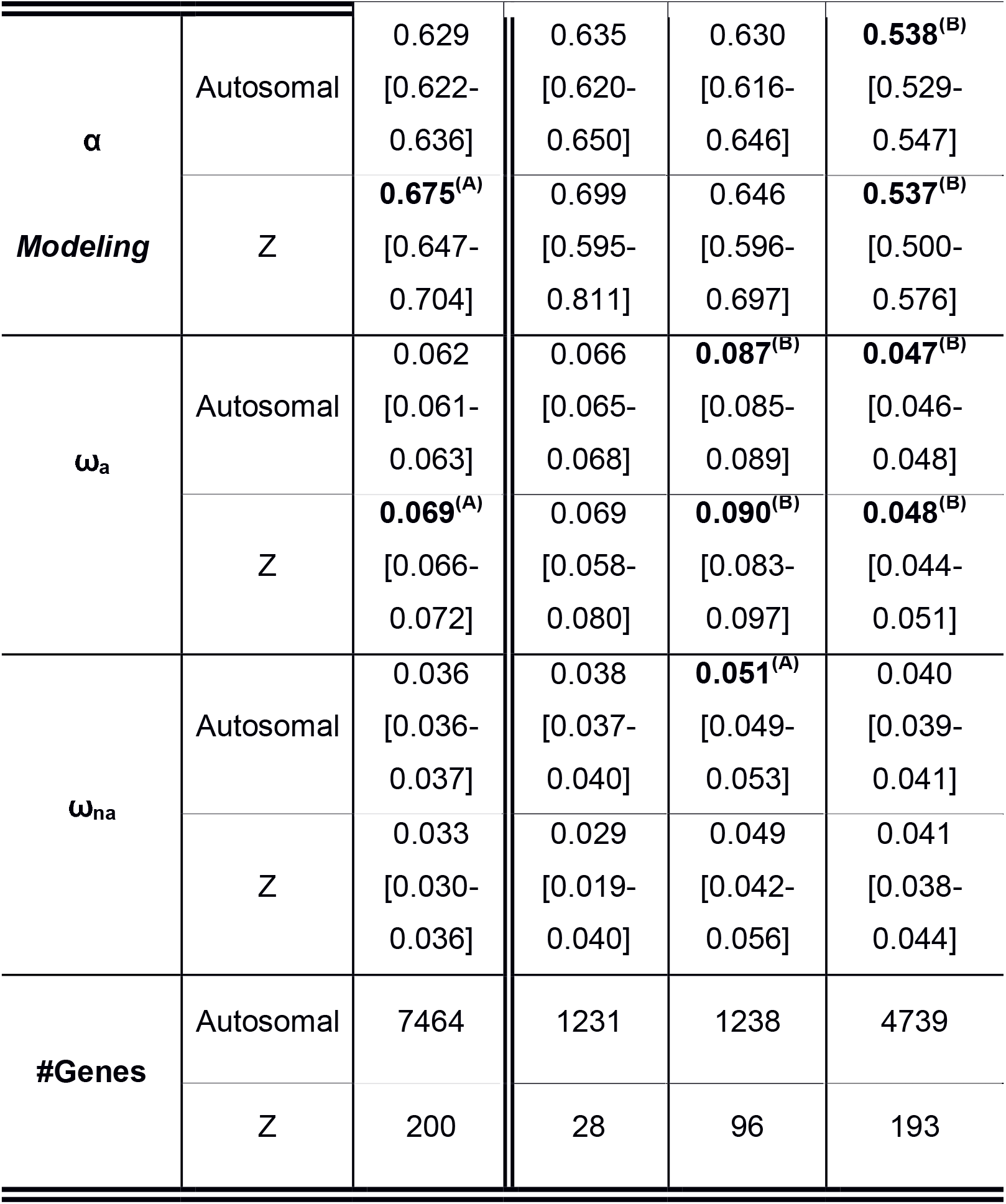
Ratios of π_n_/π_s_, dN/dS; calculations of α, ω_a_ and ω_na_ for autosomal and Z male biased, female biased and unbiased genes. π_n_/π_s_, dN/dS ratios, α, ω_a_ and ω_na_ are shown for autosomal and Z male biased, female biased and unbiased genes. Intervals represent 95% confidence intervals obtained by bootstrapping 1,000 times. **Bold**^(A)^ denotes significant values within either Z or autosomal categories. **Bold**^(B)^ denotes significant values within both Z and autosomal categories. Significance indicated separately for *All* and for sex biased expression (*female*, *male* & *unbiased*).

Finally, dS on the Z chromosome (dS_Z_=0.189; 95% CI= [0.18-0.2]) is higher than dS on the autosomes (dS_Auto_=0.162; 95% CI= [0.16-0.17]), consistent with either a: 1) male-biased mutation rate, or 2) difference in coalescence time for autosomes and Z; but does not support a fast-Z effect (Table 1).

### π_sZ_/π_sA_ diversity ratio is lower than 0.75

We next explored patterns of within-species diversity as an indicator of the strength of purifying selection. In a population at equilibrium with a 1:1 sex ratio the π_sZ_/π_sA_ diversity ratio is expected to be 0.75, but stronger purifying selection on the Z chromosome would lead to a reduction in this ratio due to background selection. The π_sZ_/π_sA_ ratio for *H. melpomene* is approximately 0.44 (Table 2), which might indicate purifying selection on the Z. However, this ratio can also be influenced by a biased sex-ratio (Vicoso & Charlesworth, 2006), differences in recombination rates (Charlesworth, 2012), sex-biased mutation rates (Vicoso & Charlesworth, 2009), or a historical reduction in population size. Recent calculations for *H. melpomene* from Panama using whole-genome short read sequencing data estimated the π_sZ_/π_sA_ diversity ratio value to be 0.611; CI= [0.570-0.653] with only weak evidence for a population bottleneck (Van Belleghem *et al*., 2018). The more pronounced reduction in diversity at synonymous sites we see here might therefore indicate enhanced background selection in genic regions of the Z chromosome.

**Table 2.**
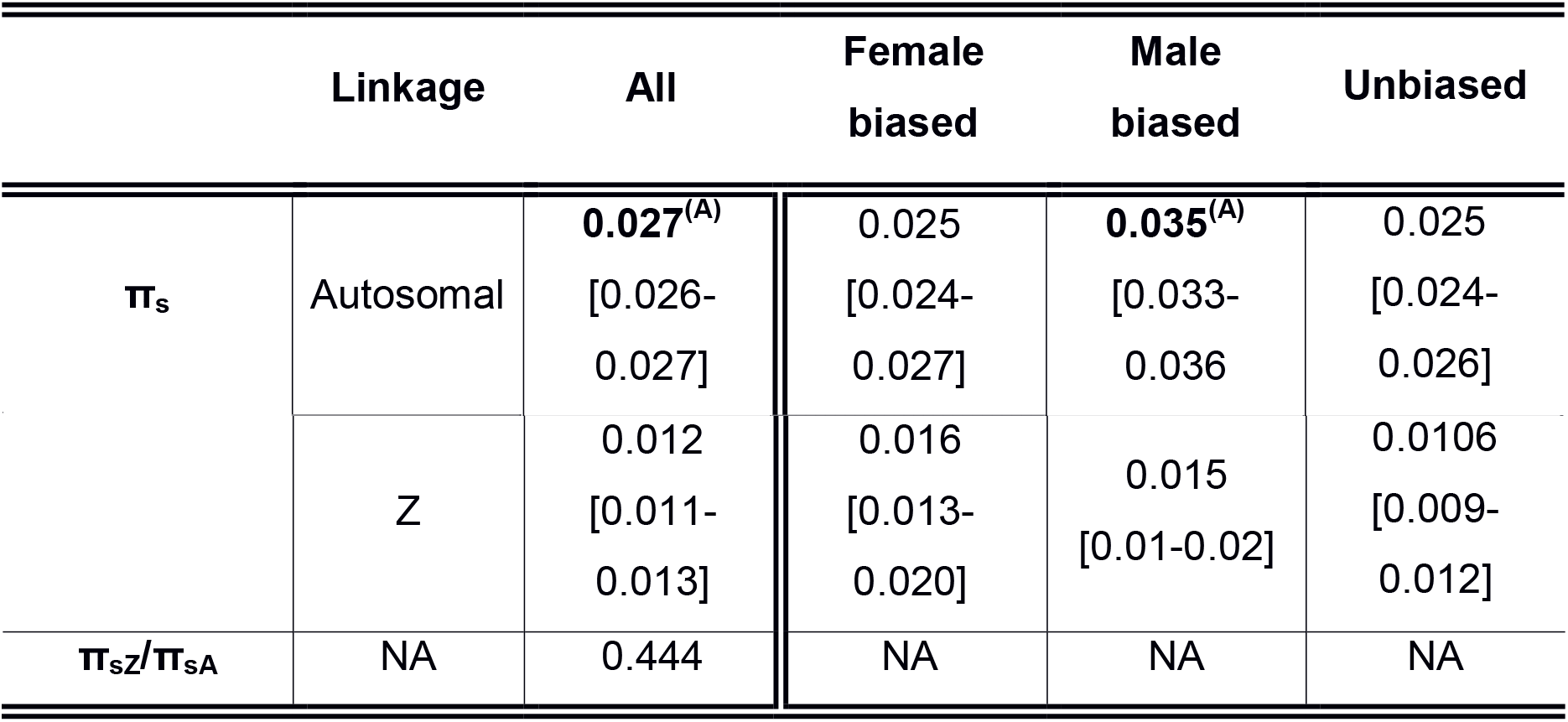
*H. melpomene* π_s_ & π_sZ_/π_sA_ ratio from pairwise alignments for Z and autosomal genes. πs calculated from pairwise alignments for Z and autosomal genes. π_sZ_/π_sA_ ratio used to estimate N_eZ_/N_eA_. Intervals represent 95% confidence intervals obtained by bootstrapping genes (1000 replicates). **Bold**^(A)^ denotes significant values within either Z or autosomal categories. Significance indicated separately for *All* and for sex biased (*female, male* & *unbiased*).

### Increased strength of purifying selection on highly expressed genes

Patterns of diversity were however strongly associated with expression levels. Using a multiple regression approach we found that functional genetic diversity, π_n_, was significantly negatively correlated with expression level for both autosomal (*P* < 0.01) consistent with increased purifying selection on highly expressed genes (Supplementary Figure S2).

**Figure 1.**
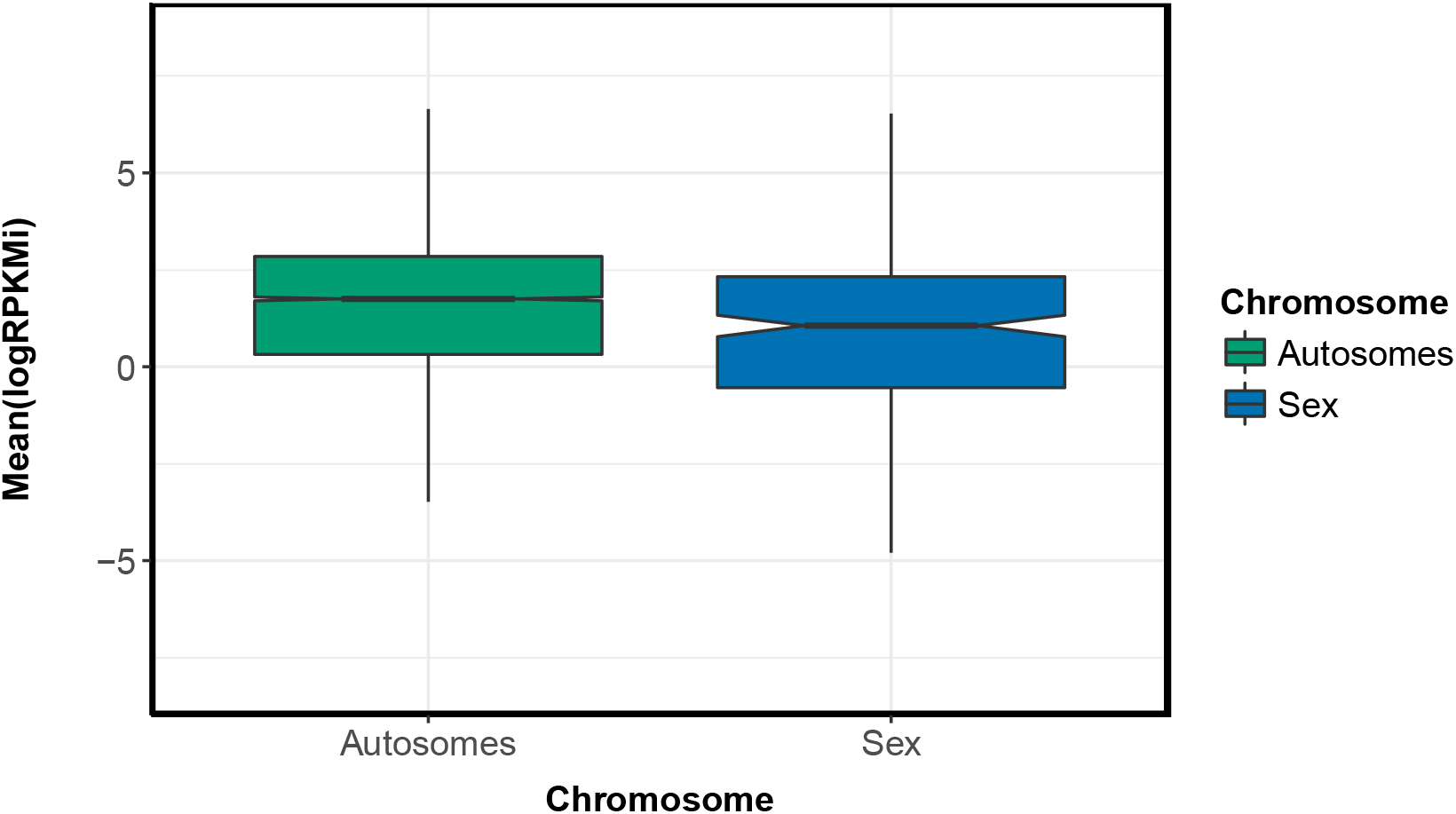
Expression level of Z and autosomal genes. Median expression level of Z genes is significantly lower than autosomal genes (*P* < 0.05). Notches on boxplot display the confidence intervals around the median.

### Z and autosomal rates of adaptive substitution: testing fast-Z adaptation

We next explored patterns of adaptive evolution using: 1) the *Classic* MK test; and 2) the *Modelling* approach which accounts for the effect of mildly deleterious mutations. We computed: 1) the proportion of adaptive non-synonymous substitutions (α) for both the *Classic* and the *Modelling* approaches; and 2) ω_a_ and ω_na_ for the *Modelling approach*. ω_a_ is the per mutation rate of adaptive substitutions and ω_na_ is the per mutation rate of non-adaptive substitutions.

There are no significant differences in α values between gene categories under the *Classic approach* (Table 1). However, using the *Modelling approach*, when all genes are considered, Z genes have a marginally but significantly higher α (α_Z_=0.675; 95% CI= [0.647-0.704]) than those that are autosomal (α_Auto_=0.629; 95% CI= [0.622-0.636]). Nonetheless, α is not significantly different between the Z chromosome and autosomes for female biased (α_Auto_=0.635; 95% CI= [0.62-0.65]; α_Z_=0.699; 95% CI= [0.595-0.811]) or male biased genes (α_Auto_=0.63; 95% CI= [0.616-0.646]; α_z_=0.646; 95% CI= [0.596-0.697]). Unbiased genes have significantly lower α values than female or male biased genes for both Z and autosomes, but within the unbiased genes there is no significant difference in α between Z (α_Z_=0.537; 95% CI= [0.5-0.567]) and autosomes (α_Auto_=0.538; 95% CI= [0.529-0.547]) (Table 1). The lack of significant differences in α between sex-biased genes is not consistent with the expectations of fast-Z adaptation, which would predict faster evolution of female biased genes due to hemizygosity compared to autosomes, but this could also reflect a lack of power to detect the signal when the total number of genes is reduced.

### Hemizygosity and the rate of adaptive substitutions

Similarly, ω_α_, the rate of adaptive substitution relative to the rate of neutral divergence, is significantly higher for Z genes (ω_αZ_=0.069; 95% CI= [0.066-0.072])as compared to autosomal genes (ω_αAuto_=0.062; 95% CI= [0.061-0.063]), consistent with the hypothesis of faster-Z evolution. However, contrary to the prediction for faster-Z evolution, ω_α_ is significantly higher for both male biased autosomal (ω_α Auto_=0.087; 95% CI= [0.085-0.089]) and Z genes (ω_α Z_=0.090; 95% CI= [0.083-0.097]) as compared to for female biased genes (ω_α Z_=0.069; 95% CI= [0.058-0.08]; ω_α Auto_=0.066; 95% CI= [0.065-0.068]). ω_α_ is significantly lower for unbiased genes as compared to male and female biased genes, both in autosomal (ω_α Auto_=0.047; 95% CI= [0.046-0.048]) and sex chromosome ω_α_ z=0.048; 95% CI= [0.044-0.051]).

There was no evidence for reduced purifying selection on the Z chromosome, as the per-mutation rate of non-adaptive substitution (ω_nα_) is lower for Z genes (ω_nαZ_ =0.033; 95% CI= [0.030-0.036] and ω_na Auto_=0.036; 95% CI= [0.036-0.037]). Female biased genes have the lowest ω_nα_ (ω_nα Auto_=0.038; 95% CI= [0.0370.040]; ω_nα Z_=0.029; 95% CI= [0.019-0.040]) compared to male biased (ω_nα Auto_=0.051; 95% CI= [0.049-0.053]; ω_nα Z_=0.049; 95% CI= [0.042-0.056]) and unbiased (ω_nα Auto_=0.04; 95% CI= [0.039-0.041]; ω_nα Z_=0.041; 95% CI= [0.038-0.044]) genes, which confirms the low π_n_/π_s_ already reported and would suggest that purifying selection is stronger in female biased genes.

### Female ovary-biased and gut-biased genes

Next we explored the expression of genes in female reproductive tissue. Overall there were a greater number of genes with gut-biased expression (#GutAuto=153) than ovary biased expression (#Ovary_Auto_=40) in the autosomes. However, there was an over-representation of Z ovary expressed genes than expected by chance (#Gut_Z_=6; #Ovary_Z_=6; chi-square test; *P* < 0.05). However, the number of genes in each category is relatively small so these tests should be treated with caution.

Of the 205 differentially expressed genes between the two tissues only 9 in the ovaries and 26 in the gut could be used to calculate dN/dS, π_n_/π_s_ and α. The other genes either do not have a 1-1 ortholog with *H. erato* or there were too many undetermined characters (gaps or Ns) to be able to estimate the parameters. Of the 35 genes for which molecular evolution statistics could be calculated, all 9 ovary biased and 25 of 26 gut biased genes are autosomal; and 1 gut biased gene maps to the Z (Gut dN/dS_Z_=0.365; Gut π_n_/π_s Z_=0.093; Gut α _Z_=0.295). We did not detect any significant differences in π_n_/π_s_; dN/dS; or α for autosomal ovary and gut-biased genes.

## Discussion

Elevated rates of coding sequence evolution on the sex chromosome relative to autosomes have been reported for several species, consistent with the theoretical prediction of fast-X evolution. Here we find evidence for enhanced rates of adaptation on the *Heliconius* Z chromosome: Z genes have a significantly higher rate of adaptive evolution when all expressed genes are considered. However, fast-X theory predicts that genes highly expressed in the hemizygous sex should be especially prone to fast-X evolution, but this prediction was not fulfilled in our data. Female-biased genes were not more fast evolving when located on the Z chromosome. The evidence for fast-Z evolution in *Heliconius* is therefore somewhat mixed.

In other taxa there is strongest support for fast-X evolution in groups with complete dosage compensation (Mank, Vicoso, *et al*., 2010; Meisel and Connallon, 2013). Theory predicts that opportunities for fast-X evolution should increase in species with somatic X-inactivation such as eutherian mammals, as there is effectively haploid expression of the sex chromosome in cells, increasing the chances of recessive beneficial mutations being fixed (Charlesworth *et al*., 1987). Groups such as Lepidoptera have been reported to have more complex patterns of sex chromosome dosage compensation. In *Heliconius* males expression of Z genes is reduced below autosomal levels, but this dosage compensation mechanism is imperfect, with males showing increased expression relative to females on the Z chromosome (Walters *et al*., 2015). However, alternative processes, such as the masculinization of the Z chromosome, may explain the apparent lack of complete dosage compensation (Gu & Walters, 2017, Huylmans *et al*. 2017). Regardless, when we compare rates of divergence and adaptation for genes with sex-biased expression, the expectations of fast-Z evolution are not clearly met. While we might expect faster rates of adaptive evolution for female-biased genes, in fact there is a weak tendency for faster rates of evolution in male-biased genes.

Although a fast-Z effect has been observed in *Bombyx mori* (Sackton *et al*., 2014), no such pattern was reported in two satyrine butterflies where the dN/dS ratio of Z genes was slightly lower than for autosomal genes (Rousselle *et al*., 2016). In *Heliconius*, although dN/dS was not significantly different between autosomal and Z genes, we did find evidence for a faster rate of adaptive substitution. Interestingly, our data also show that dS on the Z chromosome is higher than dS on the autosomes perhaps indicating a male biased mutation rate, as Z chromosomes spend more time in males than in females (Miyata *et al*., 1987). While hemizygosity is expected to expose beneficial mutations to selection and increase rates of adaptive evolution on the Z chromosome, it is also expected to increase the efficacy of purifying selection which would act to reduce evolutionary rates. It may be that the balance between these two forces differs across lepidopteran species, leading to the mixed pattern of fast-Z evolution in some taxa but not others. It is important to add, however, that the α values estimated in this study are substantially higher than α reported in Martin *et al*. (2016). Martin *et al*. (2016) estimated α using the approach developed by Messer & Petrov (2013) and, in simulations, it has been shown that it is possible that there is an overestimation of DFE-α (the method used in this study) in scenarios with strong sweeps or population expansion (Messer and Petrov, 2013).

Wright *et al*. (2015) interpreted the high dN/dS in Z genes of birds as a consequence of reduced effective population size rather than positive selection. The difference in effective population size between sex chromosomes and autosomes in female heterogametic systems is predicted to be larger than in male heterogametic systems due to higher variance of male reproductive success (Mank, Nam, *et al*., 2010). Indeed, we estimate that coding regions on the Z chromosome have an Ne 0.44 times that of autosomes. We might therefore predict a considerable reduction in the efficacy of purifying selection on butterfly Z chromosomes. This should lead to higher ω_na_ and π_n_/π_s_ ratios on the Z compared due to stronger genetic drift. However, as in satyrine butterflies (Rousselle *et al*., 2016), in *Heliconius*, ω_na_ is not higher on the Z relative to autosomes. dN/dS and π_n_/π_s_ are higher in the Z relative to autosomes in *Heliconius*, but this is not significant. This means that, in contrast to birds, the difference in the effective population size of the Z relative to autosomes is not sufficient to reduce the efficacy of purifying selection at a detectable level.

One possible explanation for this difference is the generally much higher effective population sizes of Lepidoptera, which could allow for efficient selection even on sex chromosome (Rousselle *et al*., 2016). Another is that by not using all genomic sites to estimate the π_sZ_/π_sA_ diversity ratio, we might be underestimating its true value due to a stronger effect of background selection. The latter is supported by the observation that, in a recently published paper using all genomic sites to estimate π_sZ_/π_sA_, Van Belleghem *et al*. (2018) calculated it to be 0.661 CI= [0.570-0.653]. Regardless both ours and Van Belleghem *et al*. (2018) estimates of the π_sZ_/π_sA_ diversity ratio are significantly lower than the expected 0.75 and still there is no observable reduction in the efficacy of purifying selection in *H. melpomene*.

Another factor that might counteract the fast-Z effect is adaptation from standing variation. Larger populations are more polymorphic and therefore have an increased probability of adaption from standing genetic variation. Adaptation from standing genetic variation is therefore expected to result in faster-autosome evolution, independent of the dominance of beneficial alleles (Orr and Betancourt, 2001), which would counteract the fast-Z effect. This may be especially relevant when overall population sizes are large, as in many *Heliconius* species, such that standing variation becomes a comparatively important source of adaptive variation as compared to *de novo* mutations.

As sex biased genes tend to be expressed in sex specific tissue such as the testis and the ovaries we aimed to investigate patterns of molecular evolution in ovary biased genes. Unfortunately, there are no ovary-biased genes with 1-1 orthologues between *H. melpomene* and *H. erato* that are Z. This means we could not test the effect of hemizygosity on non-sex specific and sex-specific female expression directly. The lack of 1-1 orthology may mean that these genes are rapidly evolving, and indeed autosomal ovary-expressed genes do have higher rates of adaptive evolution than gut expressed genes.

Together these results illustrate the need to study substitution rates in other ZW systems considering sex biased expression. This genome-wide analysis of polymorphism, divergence and gene expression data contributes to a growing body of literature on sex chromosome evolution in ZW systems, and reveals the complexity of the different evolutionary forces shaping transcriptome evolution in *Heliconius* and, consistent with previous work, shows limited evidence of fast-Z evolution in this taxon.

## Supplementary Tables

**Supplementary Table S1.** Sample information and statistics. *H. melpomene rosina* RNA-seq mapping statistics. Sample ID, species, tissue, stage of collection for mRNA 150bp PE directionally sequenced reads for this project. Samples mapped to *H. melpomene* genome v2.1. Walters *et al*. (2015) sample mapping statistics to *H. melpomene* genome v2.1.

| Sample | Sex | Tissue | Treatment | Library | Raw Reads |
| --- | --- | --- | --- | --- | --- |
| AP141 | Female | Gut | Old | RRB03031 | 32306468 |
| AP93 | Female | Gut | Young | RRB03025 | 34353682 |
| AP142 | Female | Gut | Old | RRB03032 | 34445514 |
| AP77 | Female | Gut | Old | RRB03030 | 32024898 |
| AP89 | Female | Gut | Old | RRB03034 | 39661158 |
| AP55 | Female | Gut | Old | RRB03035 | 35862378 |
| AP94 | Female | Gut | Young | RRB03026 | 36223403 |
| AP37 | Female | Gut | Young | RRB03029 | 31187077 |
| AP34 | Female | Gut | Young | RRB03028 | 35452206 |
| AP71 | Female | Gut | Young | RRB03024 | 33088963 |
| AP35 | Female | Gut | Young | RRB03027 | 46122142 |
| AP80 | Female | Gut | Old | RRB03033 | 40040750 |
| AP35 | Female | Ovaries | Young | RRBL00006 | 35018481 |
| AP94 | Female | Ovaries | Young | RRB02962 | 39903075 |
| AP34 | Female | Ovaries | Young | RRB02963 | 27811550 |
| AP37 | Female | Ovaries | Young | RRB02960 | 33410348 |
| AP71 | Female | Ovaries | Young | RRBL00007 | 34856038 |
| AP88 | Female | Ovaries | Young | RRBL00008 | 38486006 |
| AP93 | Female | Ovaries | Young | RRB02961 | 31497198 |
| AP55 | Female | Ovaries | Old | RRB03012 | 37934077 |
| AP77 | Female | Ovaries | Old | RRB03013 | 34322656 |
| AP80 | Female | Ovaries | Old | RRB03014 | 36157750 |
| AP89 | Female | Ovaries | Old | RRB03015 | 34318423 |
| AP141 | Female | Ovaries | Old | RRB03016 | 33844256 |
| AP142 | Female | Ovaries | Old | RRB03017 | 35328097 |
| R20 | Female | Abdomen | Young | NA | NA |
| R29 | Female | Abdomen | Young | NA | NA |
| R06 | Male | Abdomen | Young | NA | NA |
| R32 | Male | Abdomen | Young | NA | NA |
| R34 | Male | Abdomen | Young | NA | NA |
| R07 | Female | Abdomen | Young | NA | NA |
| R33 | Male | Abdomen | Young | NA | NA |
| R05 | Female | Abdomen | Young | NA | NA |
| R21 | Male | Abdomen | Young | NA | NA |
| R28 | Female | Abdomen | Young | NA | NA |

**Supplementary Table S1. Sample information and statistics (cont.)**
| **Sample** | **Raw Base (G)** | **Error Rate (%)** | **Q20 (%)** | **Q30 (%)** | **GC content (%)** |
| --- | --- | --- | --- | --- | --- |
| AP141 | 9.69 | 0.01 | 98.25 | 95.7 | 41.52 |
| AP93 | 10.31 | 0.01 | 97.38 | 93.98 | 40.89 |
| AP142 | 10.33 | 0.01 | 98.25 | 95.75 | 41.5 |
| AP77 | 9.61 | 0.01 | 98.23 | 95.69 | 42.14 |
| AP89 | 11.9 | 0.01 | 98.11 | 95.53 | 42.53 |
| AP55 | 10.76 | 0.01 | 98.15 | 95.6 | 41.52 |
| AP94 | 10.87 | 0.01 | 97.94 | 95.03 | 41.41 |
| AP37 | 9.36 | 0.01 | 98.52 | 96.42 | 42.95 |
| AP34 | 10.64 | 0.01 | 98.53 | 96.4 | 41.49 |
| AP71 | 9.93 | 0.01 | 98.12 | 95.39 | 40.62 |
| AP35 | 13.84 | 0.01 | 97.84 | 94.81 | 40.47 |
| AP80 | 12.01 | 0.01 | 98.19 | 95.62 | 42 |
| AP35 | 10.51 | 0.01 | 98.53 | 96.41 | 41.08 |
| AP94 | 11.97 | 0.01 | 98.16 | 95.19 | 41.51 |
| AP34 | 8.34 | 0.01 | 98.2 | 95.27 | 41.42 |
| AP37 | 10.02 | 0.01 | 98.49 | 95.88 | 42.15 |
| AP71 | 10.46 | 0.01 | 98.42 | 96.19 | 41.36 |
| AP88 | 11.55 | 0.01 | 98.53 | 96.37 | 42.71 |
| AP93 | 9.45 | 0.01 | 98.38 | 95.69 | 41.24 |
| AP55 | 11.38 | 0.01 | 98.17 | 95.51 | 39.76 |
| AP77 | 10.3 | 0.01 | 98.28 | 95.6 | 40.46 |
| AP80 | 10.85 | 0.01 | 97.97 | 95.06 | 40.28 |
| AP89 | 10.3 | 0.01 | 98.04 | 95.16 | 41.51 |
| AP141 | 10.15 | 0.01 | 97.88 | 94.83 | 39.58 |
| AP142 | 10.6 | 0.01 | 98 | 95.09 | 39.69 |
| R20 | NA | NA | NA | NA | NA |
| R29 | NA | NA | NA | NA | NA |
| R06 | NA | NA | NA | NA | NA |
| R32 | NA | NA | NA | NA | NA |
| R34 | NA | NA | NA | NA | NA |
| R07 | NA | NA | NA | NA | NA |
| R33 | NA | NA | NA | NA | NA |
| R05 | NA | NA | NA | NA | NA |
| R21 | NA | NA | NA | NA | NA |
| R28 | NA | NA | NA | NA | NA |

**Supplementary Table S1. Sample information and statistics**
| **Sample** | **Mapped Reads(%)** | **Properly Paired(%)** |
| --- | --- | --- |
| AP141 | 83.40 | 78.16 |
| AP93 | 75.27 | 69.38 |
| AP142 | 79.84 | 74.32 |
| AP77 | 79.71 | 74.03 |
| AP89 | 73.41 | 67.05 |
| AP55 | 77.95 | 72.08 |
| AP94 | 62.79 | 58.07 |
| AP37 | 76.98 | 72.13 |
| AP34 | 81.84 | 76.85 |
| AP71 | 79.86 | 74.15 |
| AP35 | 76.68 | 71.58 |
| AP80 | 75.17 | 69.39 |
| AP35 | 76.76 | 71.96 |
| AP94 | 82.03 | 77.30 |
| AP34 | 86.45 | 81.59 |
| AP37 | 83.17 | 78.42 |
| AP71 | 85.70 | 80.81 |
| AP88 | 86.99 | 82.85 |
| AP93 | 82.80 | 78.53 |
| AP55 | 85.29 | 80.51 |
| AP77 | 87.83 | 83.17 |
| AP80 | 86.56 | 81.74 |
| AP89 | 85.34 | 80.68 |
| AP141 | 88.09 | 83.44 |
| AP142 | 87.19 | 82.62 |
| R20 | 38.20 | 34.46 |
| R29 | 31.92 | 29.07 |
| R06 | 60.95 | 53.47 |
| R32 | 85.00 | 77.32 |
| R34 | 29.08 | 26.05 |
| R07 | 83.25 | 76.15 |
| R33 | 35.84 | 30.56 |
| R05 | 71.44 | 64.81 |
| R21 | 38.06 | 31.36 |
| R28 | 61.37 | 54.10 |

**Supplementary Table S2.** Orthologue prediction improvement with Hmel2.5 annotation. Statistics on orthologue prediction between *H. melpomene* v2 annotation and *H. erato* annotation; and on orthologue prediction between *H. melpomene* v2.5 annotation and *H. erato* annotation.

| **Statistics** | **Hmel2** | **Hmel2.5** |
| --- | --- | --- |
| # genes | 33137 | 41779 |
| # genes in orthogroups | 24274 | 29698 |
| # unassigned genes | 8863 | 12081 |
| % genes in orthogroups | 73.3 | 71.1 |
| % unassigned genes | 26.7 | 28.9 |
| # orthogroups | 9320 | 11062 |
| # species-specific orthogroups | 15 | 18 |
| # genes in species-specific orthogroups | 56 | 105 |
| % genes in species-specific orthogroups | 0.2 | 0.3 |
| Mean orthogroup size | 2.6 | 2.7 |
| Median orthogroup size | 2.0 | 2.0 |
| G50 assigned genes | 2 | 2 |
| G50 all genes | 2 | 2 |
| O50 assigned genes | 3252 | 3638 |
| O50 all genes | 5468 | 6658 |
| # of orthogroups with all species present | 9305 | 11044 |
| # of single-copy orthogroups | 6846 | 8095 |

**Supplementary Table S3.** Mean and median read depth (RD) for resequenced whole-genome *H. melpomene* samples (van Belleghem *et al*., 2018) *H. melpomene* resequenced samples mapped to Hmel2 genome using BWA-MEM

| Sample | Species | Sex | Location | Mean RD | Median |
| --- | --- | --- | --- | --- | --- |
|  |  |  |  |  | RD |
| CAM000531 | H. m. rosina | Male | 9°87'N 7°96'W | 30.59 | 29 |
| CAM000533 | H. m. rosina | Male | 9°87'N 7°96'W | 28.83 | 21 |
| CAM000546 | H. m. rosina | Male | 9°87'N 7°96'W | 27.47 | 26 |
| CAM001841 | H. m. rosina | Male | 9°87'N 7°96'W | 28 | 28 |
| CAM001880 | H. m. rosina | Male | 9°87'N 7°96'W | 22.76 | 23 |
| CAM002045 | H. m. rosina | Male | 9°87'N 7°96'W | 25.7 | 26 |
| CAM002059 | H. m. rosina | Male | 9°87'N 7°96'W | 36.77 | 32 |
| CAM002071 | H. m. rosina | Male | 9°87'N 7°96'W | 26.43 | 21 |
| CAM002519 | H. m. rosina | Male | 9°87'N 7°96'W | 26.68 | 22 |
| CAM002552 | H. m. rosina | Male | 9°87'N 7°96'W | 26.83 | 22 |

## Supplementary Figures

**Supplementary Figure S1.**
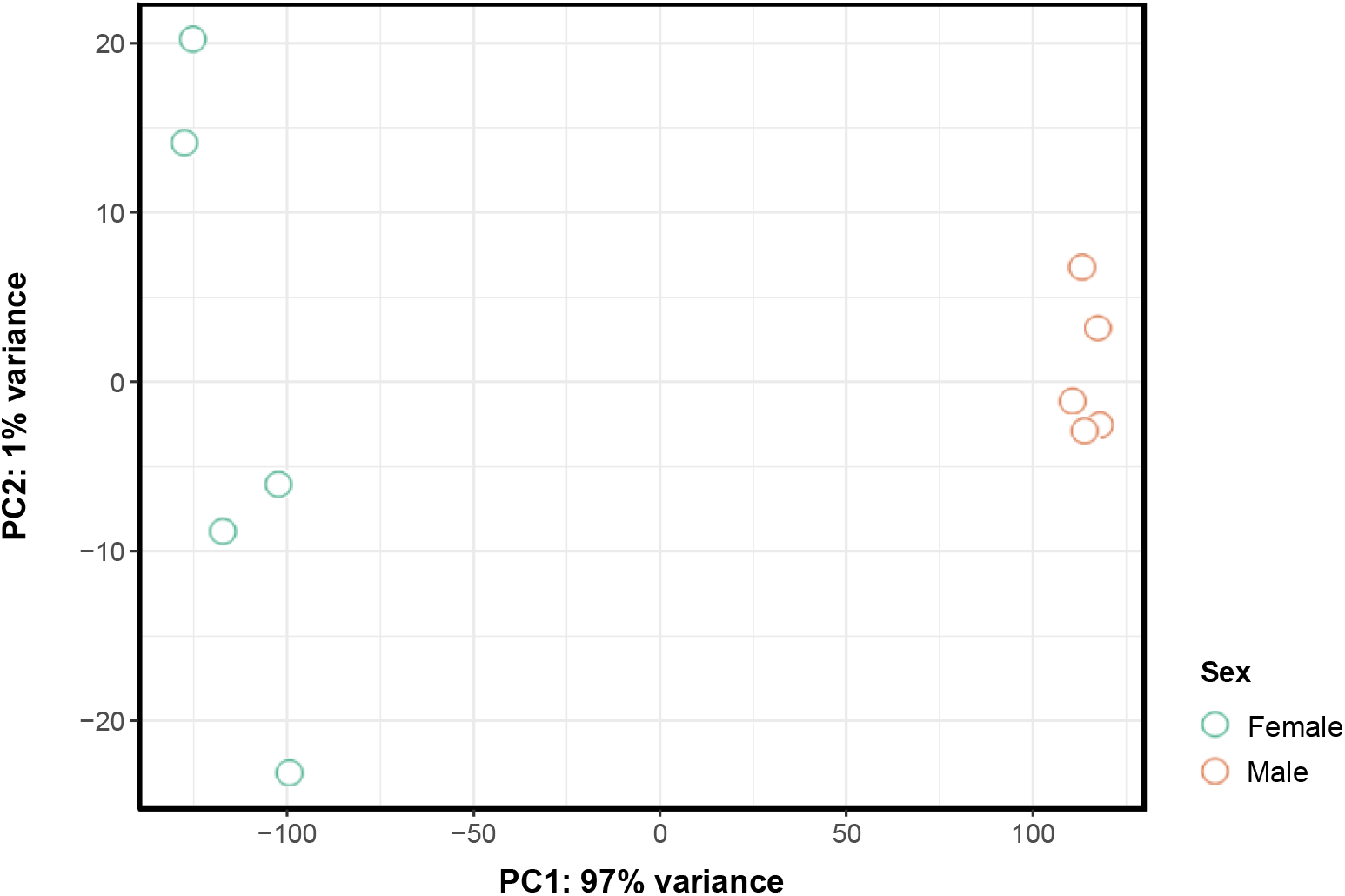
Principal component analysis of gene expression profiles for the 10 whole abdomen male and female samples. PCA of the abdomen transformed gene expression count data to the log2 scale (DESeq2, rlog(blind=FALSE)). rlog transformed data minimises differences between samples for rows with small counts and normalizes with respect to library size.

**Supplementary Figure S2.**
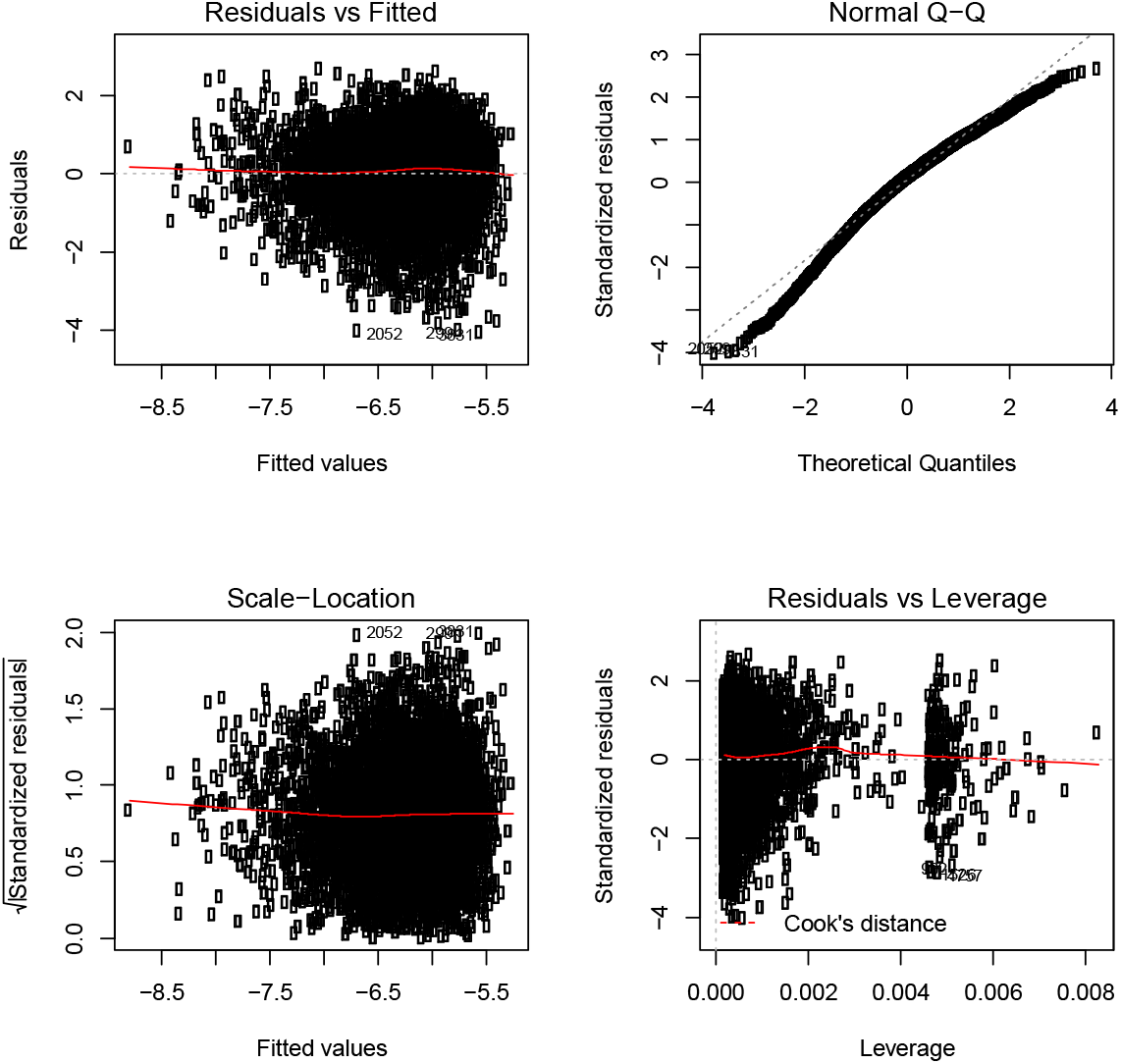
π_n_ is negatively correlated to expression level. Multiple regression approach shows that π_n_ was significantly negatively correlated to expression level – autosomal (*P* < 0.001) and Z genes (*P* < 0.01). **A.** Plotted *Residuals vs Fitted* shows spread residuals around the horizontal line without distinct patterns. *Normal Q-Q* follow a straight line with residuals well lined. The *Scale-Location* plot shows residuals spread equally around range of predictors. There is equal variance or homoscedasticity. *Residuals vs Leverage* plot does not identify any influential outliers in the linear regression analysis. **B.** Regression coefficient table. Relationship between π_n_ and expression.

**Supplementary Figure S3.**
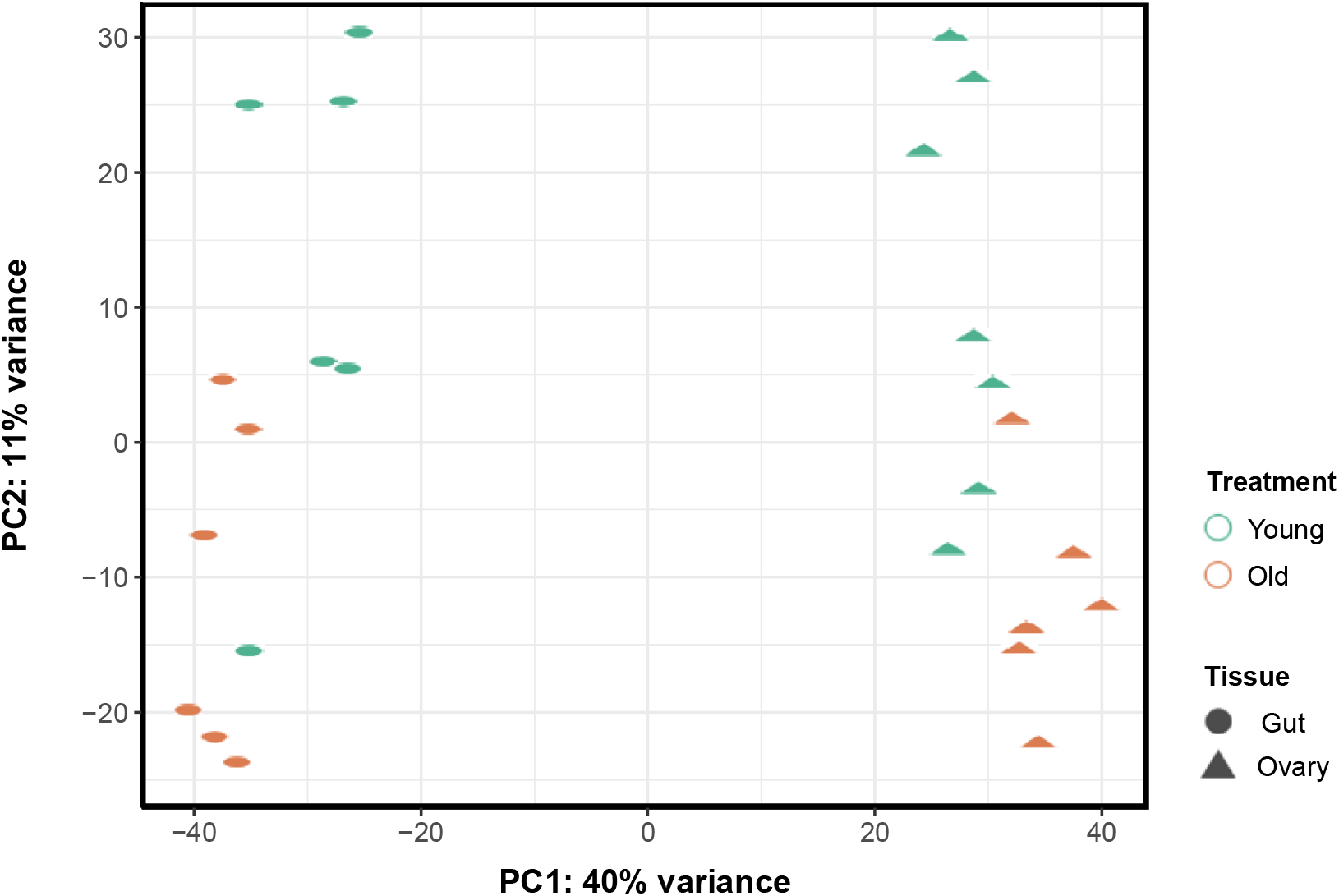
Principal component analysis of gene expression profiles of *H. melpomene* females for 13 ovary samples and 12 gut samples at two different time points. PCA of the female ovary and gut transformed gene expression count data to the log2 scale (DESeq2, rlog(blind=FALSE)). rlog transformed data minimises differences between samples for rows with small counts and normalizes with respect to library size.

**Supplementary Figure S4.**
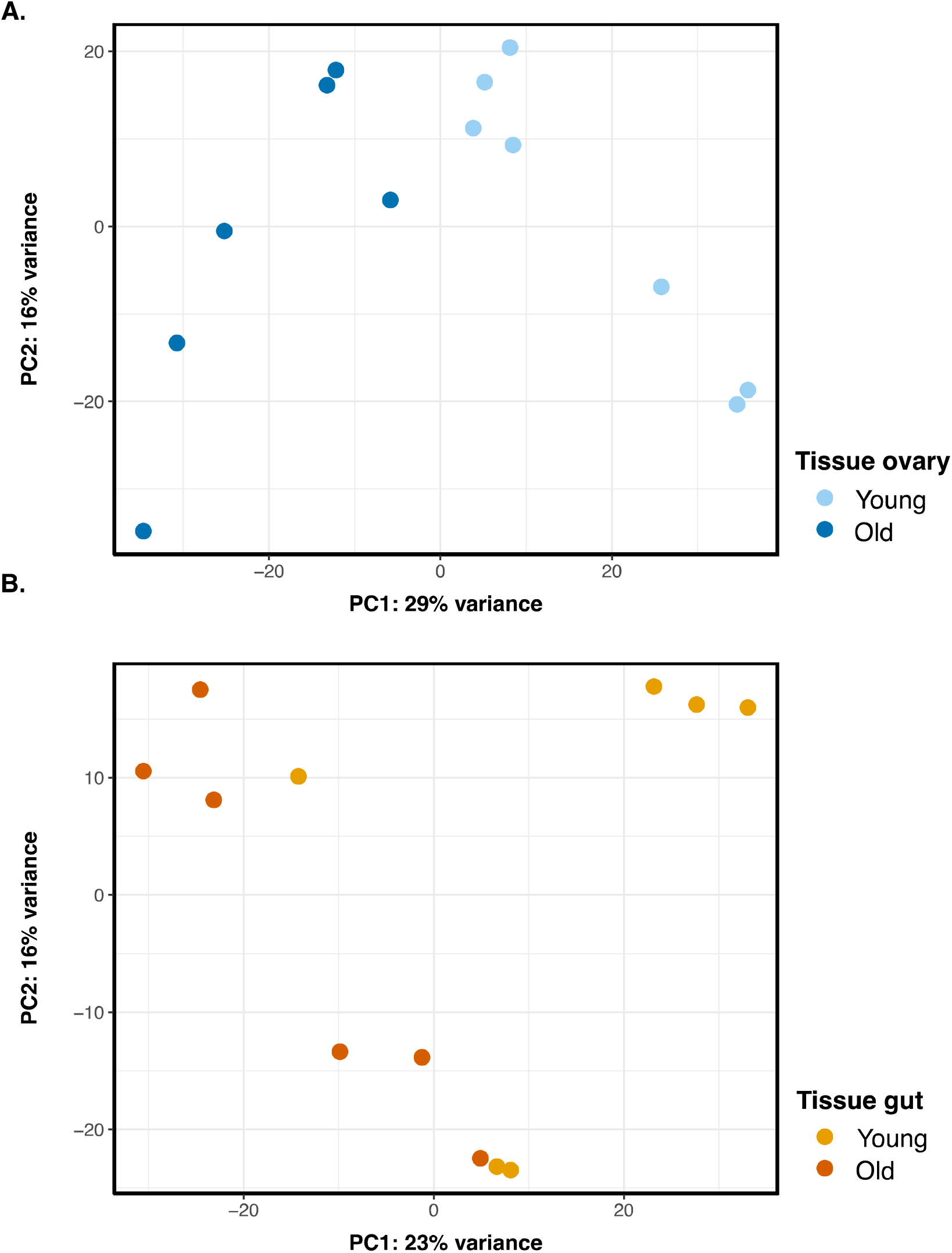
Principal component analysis of gene expression profiles of *H. melpomene* females for 13 ovary samples and 12 gut samples at two different time points separated by tissue type. PCA of the female ovary and gut transformed gene expression count data to the log2 scale (DESeq2, rlog(blind=FALSE)) separated by tissue. rlog transformed data minimises differences between samples for rows with small counts and normalizes with respect to library size. **A.** 45% of the variance is explained by PC1 and PC2. PC1 separates young ovary tissue from old ovary tissue and explains 29% of the variance. All the samples cluster by age. **B.** 39% of the total variance is explained by PC1 and PC2. PC1 separates young gut tissue from old gut tissue and explains 23% of the variance. The samples cluster less tightly by age than ovary expression.

